# The influence of X chromosome variants on trait neuroticism

**DOI:** 10.1101/401166

**Authors:** Michelle Luciano, Gail Davies, Kim M Summers, W David Hill, Caroline Hayward, David C Liewald, David J Porteous, Catharine R. Gale, Andrew M McIntosh, Ian J Deary

**Author notes:** Corresponding author, Department of Psychology, The University of Edinburgh, 7 George Square, EH8 9JZ, Edinburgh, UK.

## Abstract

Autosomal variants have successfully been associated with trait neuroticism in genome-wide analysis of adequately-powered samples. But such studies have so far excluded the X chromosome from analysis. Here, we report genetic association analyses of X chromosome and XY pseudoautosomal single nucleotide polymorphisms (SNPs) and trait neuroticism using UK Biobank samples (N = 405,274). Significant association was found with neuroticism on the X chromosome for 204 markers found within three independent loci (a further 783 were suggestive). Most of these significant neuroticism-related X chromosome variants were located in intergenic regions (n = 713). Involvement of *HS6ST2*, which has been previously associated with sociability behaviour in the dog, was supported by single SNP and gene-based tests. We found that the amino acid and nucleotide sequences are highly conserved between dogs and humans. From the suggestive X chromosome variants, there were 19 nearby genes which could be linked to gene ontology information. Molecular function was primarily related to binding and catalytic activity; notable biological processes were cellular and metabolic, and nucleic acid binding and transcription factor protein classes were most commonly involved. X-variant heritability of neuroticism was estimated at 0.34% (SE = 0.07). A polygenic X-variant score created in an independent sample (maximum N ≈ 7300) did not predict significant variance in neuroticism, psychological distress, or depressive disorder. We conclude that the X chromosome harbours significant variants influencing neuroticism, and might prove important for other quantitative traits and complex disorders.

---

Neuroticism is a dispositional personality trait tapping negative emotion, like experience of anxiety, mood swings, and negative affect. It is a predictor of a range of psychiatric traits and disorders ^1,2^. Our understanding of the genetic influences on people’s differences in neuroticism has increased substantially with the availability of large biological resources like UK Biobank. In a genome-wide autosomal analysis of neuroticism in UK Biobank, Luciano and colleagues ^3^ found 116 independent single nucleotide polymorphisms (SNPs) associated with neuroticism and replicated 15 of them in a large independent sample, 23andMe. These variants and the biological pathways that were identified provided some understanding about the biological processes and molecular mechanisms of importance, for example, neural genesis and differentiation pathways. There was genetic overlap between neuroticism and depressive disorder and anti-depressant drug targets. Large-scale association analysis of neuroticism and the genetic variants on the sex chromosomes has not yet been performed; therefore, our current understanding of the genetic factors that contribute to neuroticism is incomplete. The analysis of X chromosome variation (including pseudoautosomal variation) and neuroticism, in UK Biobank, is the focus of the current study.

In humans, phenotypic sex is determined by the X and Y sex chromosomes, with females having two X chromosomes and males having a single X chromosome and a single Y chromosome. To prevent overexpression of genes on the X chromosome in females, one X is randomly inactivated during development so that females have only one active X in each cell. This has implications for genetic mutations in genes located on the X chromosome and can result in phenotypes having a different prevalence between the sexes ^4^. The X chromosome has a role in the development of the human brain ^5^. Because men and women differ in their mean level of some but not all personality traits ^6^, consideration of the X chromosome is important in understanding the genetic architecture of these traits.

Neuroticism scores are generally higher in women than men ^6,7^, and neuroticism-related mood disorders such as anxiety and depression show greater prevalence in women than in men ^8^. Findings from twin studies suggested that the genetic influences on neuroticism might be larger for women than men ^9,10^. In 6,828 twin brothers and 8,104 twin sisters from Finland aged 24 to 53 years, heritability of neuroticism (9-item Eysenck scale) differed significantly between men (.31) and women (.42) ^11^. Opposite-sex twins were not sampled, precluding analysis of qualitative sex limitation, that is, where trait variation is influenced by different factors between the sexes. An integrated analysis of this Finnish sample with twins and extended family members from Australia (n = 7,532) and the US (n = 20,554) showed that the same genes influenced neuroticism in both sexes, but that non-additive genetic effects were significantly larger in men than women (21.9% versus 13.1%; additive genetic variance of 15.7% versus 38.2%) ^12^. In 48,850 Australian and US individuals from extended twin families, broad heritability of neuroticism was slightly higher in women (41%) than men (35%) ^13^.

An alternative method of estimating trait heritability in unrelated individuals using observed genotype data found broadly consistent results to the pedigree-based analyses. In a population sample (5,016 male, 6,945 female), the heritability of neuroticism, estimated by common autosomal genetic variants, did not differ between sexes ^14^. However, point estimates of heritability were higher for men than women (.16, SE .07 versus .06, SE .05; SNP x sex interaction effect P-value = .08). In this same study, analysis of the X chromosome was not-significant, that is, X variants did not contribute to the heritability of neuroticism. Here, we reconsider the contribution of X chromosome variance in neuroticism based on an approximate 34-fold increased sample size.

With regard to the identification of X chromosome loci influencing neuroticism, the largest genetic linkage study of neuroticism, in around 5,000 sibling pairs, did not show evidence of linked QTLs on the X chromosome ^15^. Genome-wide association studies (GWAS), which are able to localise gene variant effects, typically focus on the autosomes; several small GWAS of neuroticism (e.g., N ranging ≈2200 to ≈4000) have analysed the X chromosome with null results ^16,17^. The present GWAS comprises over 400,000 UK Biobank participants to test whether X chromosome variants are associated with neuroticism scores. Further, we investigate the XY pseudoautosomal region (the ends of each arm of the chromosome that are common to both sex chromosomes, escape X-inactivation and participate in recombination) ^18^. Results will be of special value to research on mood disorders, which are genetically highly correlated with neuroticism ^19^.

## Materials and Methods

Neuroticism was measured in UK Biobank, an open resource including more than 500,000 adults aged 40 to 69 years residing in the UK (**http://www.ukbiobank.ac.uk**), with data collected from 2006 through to 2010 ^20^. We have previously published genome-wide autosomal results on neuroticism in unrelated individuals from this cohort (N = 329,824) ^3^; here we include related individuals and focus on the X chromosome (N = 405,274) ^21^. X chromosome and XY pseudoautosomal genetic variants were analysed for their association with neuroticism, measured as the summed score of 12 items from the Eysenck Personality Questionnaire-Revised Short Form (EPQ-R-SF) ^22^. Information on the imputation of missing data and its distribution can be found in our previous report ^3^. Neuroticism scores were residualised for age, assessment centre, genotype batch, array, and 40 genetic principal components. 408,091 participants (45.94% male) of European descent had high-quality genotyping imputed to the UK10K haplotype, 1000 Genomes Phase 3, and Haplotype Reference Consortium panels. Quality control of this dataset was performed as described by Bycroft and colleagues ^21^. Further quality control filters were applied before analysis: individuals were removed sequentially based on non-British ancestry, high missingness, high relatedness (samples which have more than 10 putative third-degree relatives), and gender mismatch. All bi-allelic variants with a minor allele frequency ≥ 0.000009 and an information/imputation quality score of ≥ 0.1 were analysed.

Regression models of neuroticism on genetic variants (accounting for genotype uncertainty) were run separately for men and women, specifying male genotypes as 0/2 and female genotypes as 0/1/2 which assumes random X inactivation. Association analyses were performed using BOLT-LMM software, using a linear mixed model ^23^. Meta-analysis of the association results of men (n=186,015) and women (n=219,259) allowed increased statistical power and by incorporating heterogeneity tests we could investigate differences in the magnitude of variant effects between the sexes ^24^. On the X chromosome, 1,686,693 SNPs were meta-analysed. Pseudoautosomal regions were treated as autosomes and analysed under an additive model (n = 40,764 SNPs).

### Downstream mapping of X chromosome association signals

FUnctional Mapping and Annotation of genetic associations (FUMA) was used to interrogate the association results, including identification of independent association signals, gene-based analysis, functional annotation, and gene expression ^25^. Independent significant SNPs were those with a P ≤5×10^−8^ and not in linkage disequilibrium (LD; *r*^2^<.6) with other significant SNPs. Tagging SNPs were those with MAF?≥.0005 and in LD (*r*^2^<.6) with one or more of the independent significant SNPs; they were from the 1000G reference panel and not necessarily included in the present association analysis. Any associated loci within 250kb of each other were considered a single locus. Lead SNPs were those independent significant SNPs that were even less closely related (LD *r*^2^<.1). Gene-based analysis is performed in FUMA using MAGMA ^26^. Using the default settings, SNPs were allocated to genes according to the NCBI 37.3 build positions without extending the boundary beyond the gene. LD was taken into account with reference to the European panel of the 1000G data (phase 1, release 3). Gene-based statistics were available for 762 genes, but given that autosomal genes have been previously tested in this sample ^3^, a genome-wide threshold for significance was adopted α < 2.65×10^−6^ (.05/18,842). Expression quantitative trait loci (eQTL) and functional annotation in FUMA relies on the publicly available Genotype-Tissue Expression Portal (GTEx) and Regulome DB databases.

### Estimation of X chromosome SNP-based heritability

A genetic relationship matrix (GRM) was created using the autosomal SNPs; a relatedness cut-off of 0.05 was applied to this GRM. The resulting set of unrelated individuals were used in the following GCTA-GREML analyses ^27^. We created an X chromosome GRM using the previously-defined set of unrelated individuals (N = 313,467). This GRM was then parametrized under three assumptions of dosage compensation ^28^: 1. Equal X-linked variance for males and females; 2. No dosage compensation, meaning that each allele has a similar effect in both males and females, but, as both X chromosomes are active for females, and only one in males, there is twice the amount of genetic variance linked to the X chromosome in females; 3. Full dosage compensation, i.e., that one of the X chromosomes is completely inactive for females, meaning that in females each allele has only half the effect of an allele in a male. Under full dosage compensation for females, genetic variance on the X chromosome will be around half of what will be observed in males. To estimate the <u>X</u> proportion of variance explained by X chromosome common SNPs (<u>h</u> <u>^2^</u>), we fitted the parameterized GRMs for the X chromosome in a mixed linear model whilst simultaneously estimating the proportion of variance explained by all common autosomal SNPs (*h*^*2*^). These analyses were performed on a random sub-sample of ≈150,000 individuals due to computing resource capacity.

### Polygenic Prediction into Generation Scotland

Neuroticism polygenic X chromosome prediction of neuroticism, psychological distress, and depressive disorder was carried out in Generation Scotland (GS) ^29^. Polygenic scores were calculated in PLINK ^30^ based on the female and male SNP association results for neuroticism in UK Biobank. Prediction was carried out in unrelated individuals (n ranged 3,908-4,189 for women; 3,004-3,179 for men). Only SNPs in linkage equilibrium (r^2^<.10 within a 250kb window) were included in the polygenic score. Individuals from GS who participated in UK Biobank were excluded (n= 290 of those who were unrelated). P-value thresholds of .01, .05, .1, .5 and 1 for the X chromosome association results were used to create 5 polygenic scores. Polygenic scores for each threshold were based on 163 (female)/166 (male) SNPs, 527/514 SNPs, 893/879 SNPs, 2788/2786 SNPs, and 4122/4137 SNPs, respectively. The associations between the polygenic score and neuroticism/psychological distress in GS were tested using linear regression, controlling for age and the first 10 genetic principal components. Depression status was analysed by logistic regression with the same covariate controls. To correct for multiple testing across threshold and trait, a false discovery rate (FDR) method was used ^31^.

### Phylogenetic analysis

A phylogenetic tree for the *HS6ST2* gene was retrieved using the Ensembl database *Gene tree* function (Ensembl release 93, accessed July 2018; http://www.ensembl.org). All nodes except those containing the human and dog genes were collapsed for ease of visualisation. Predicted protein sequences for HS6ST2 from a range of species were downloaded from Ensembl. The longest available predicted protein sequence was captured for each species. The sequences were aligned using Clustal Omega (https://www.ebi.ac.uk/Tools/msa/clustalo/) with the default parameters.

## Results

X chromosome and XY pseudoautosomal meta-analysis association results showed evidence of genomic inflation (quantile-quantile plots in Supplementary Figures 1 and 2), with respective lambdas of 1.07 and 1.06. There were 204 SNPs that reached genome-wide significance (P < 5×10^−8^) for the X chromosome, but none for the XY pseudoautosomal region. At a suggestive level (P < 1×10^−5^), a further 783 X chromosome variants and 3 XY pseudoautosomal variants were identified (Supplementary Tables 1 and 2). X and XY pseudoautosomal association P-values are depicted in Figure 1; Supplementary Table 3 shows annotation of the significant 1000G SNPs. Sex-stratified association results can be found in Supplementary Figures 3 and 4; none were significant. Among the significant X chromosome SNPs from the meta-analysis, there were three distinct loci: Xp21.3 (183.8kb, containing 65 SNPs), Xq25 (295.2kb, 309 SNPs), and Xq26.2 (122.4kb, 41 SNPs). Regional association plots for these loci are shown in Supplementary Figures 5-7. The respective lead SNPs (significant and with r^2^ <.1) within these regions were rs6630665 (P = 1.023×10^−8^), rs177010 (P = 2.594×10^−11^), and rs5977754 (P = 2.521×10^−8^). 397 SNPs were intergenic and 48 intronic. One protein-coding gene was mapped to the associated part of Xq26.2 (131817931-131940379bp): *HS6ST2* (Heparan Sulfate 6-O-Sulfotransferase 2); there were no protein coding genes in the loci identified within Xp21.3 and Xq25.

**Figure 1.**
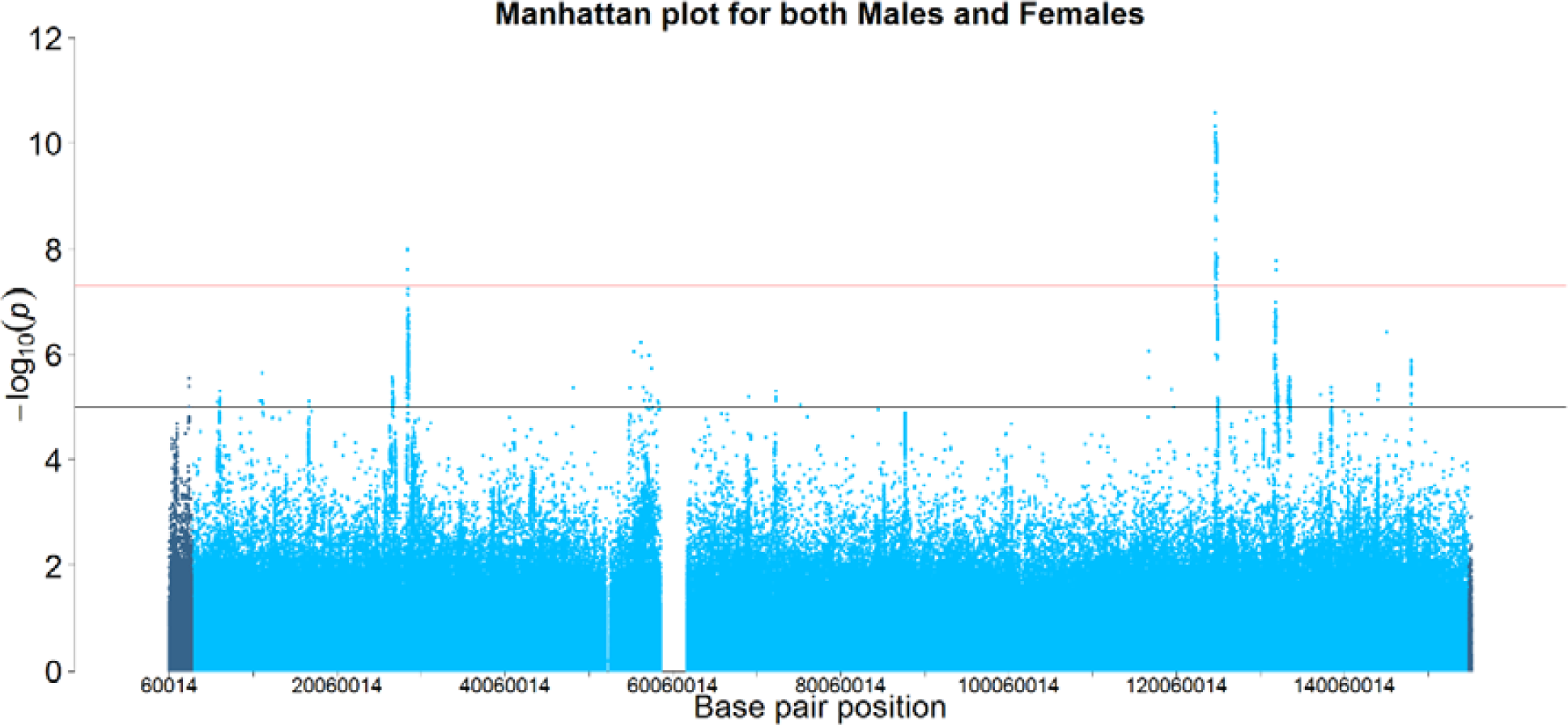
X Chromosome association results for neuroticism in UK Biobank (N = 405,274). XY pseudoautosomal SNPs are in darker blue.

**Figure 2.**
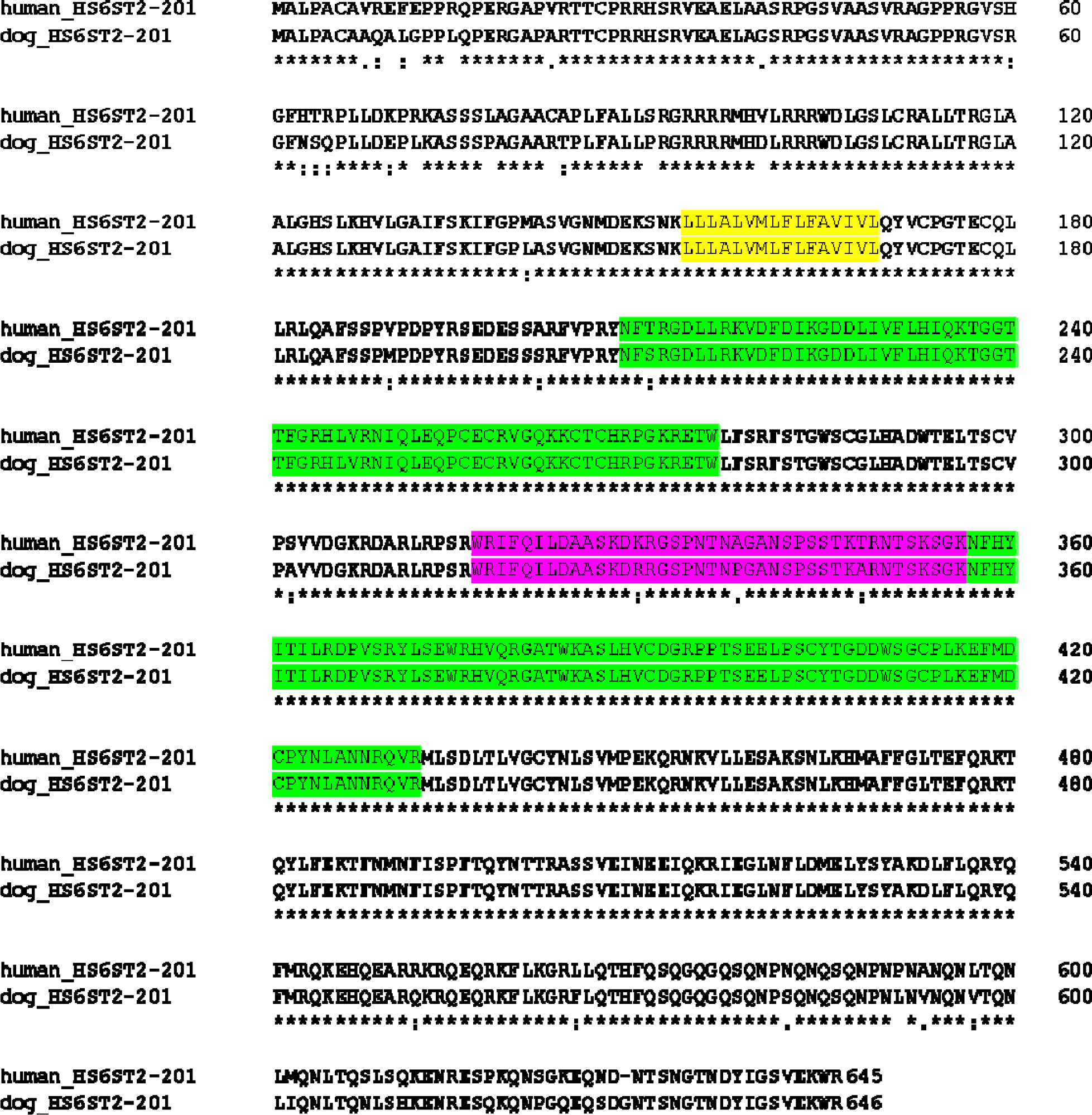
Allsnment of human and canine *HS6ST2* senes. The alignment was created usins Clustal Omesa. The human sequence accession ID is ENSP00000429473/1-645and the canine sequence accession ID is ENSCAFP00000038323/1/646. The putative transmembrane domain is hishlighted in yellow; the two P-loop containing NTP hydrolase domains are hishlighted in sreen and the resion removed by alternate splicing that produces the short form is highlishted in pink (information derived from Habuchi et al 2003 and Ensembl). Asterisks indicate positions with a conserved amino acid; colons indicate conservation of amino acids with hishlysimilar properties; periods indicate conservation of amino acids with weakly similar properties.

None of the four independent SNPs (significant and with r^2^ <.6 rather than r^2^ <.1 for lead SNPs; 2 in Xq25) were expression quantitative trait loci (eQTLs; Supplementary Table 3). Regulome DB scores (http://www.regulomedb.org/) were available for two of the independent SNPs (rs6630665, rs5977754), which showed minimal DNA binding evidence, and thus, unlikely to have a regulatory role (Supplementary Table 3). The Protein Analysis Through Evolutionary Relationships (PANTHER) classification system ^32^ was used to describe the genes located near our suggestive X chromosome variants. Of the 30 genes identified, 19 could be mapped to different biological (23 hits), cellular (11), molecular (13), and protein (17) classes (Supplementary Figures 5-8). Consistent with classification of the prominent genes from autosomal analysis ^3^, metabolic and cellular processes were the primary biological classes, nucleic acid binding and transcription factor were among the most important protein classes, and binding and catalytic activity were the principal molecular functions.

Gene-based tests for the X chromosome showed that 7 genes were significantly associated with neuroticism (Supplementary Table 4 shows all gene results). They included: *HS6ST2* (containing 1190 SNPs)*, IL1RAPL1* (6670)*, HPRT1* (226)*, PHF6* (187)*, DMRTC1* (50)*, CTPS2* (528), and *NLGN4X* (2139). Only one—*HS6ST2*—showed single SNPs attaining significance. For the XY pseudoautosomal region, the *XG* gene (including 45 SNPs) was mapped and was not significant (P = .5).

Genome Complex Trait Analysis (GCTA) ^28^ SNP heritability estimates did not differ significantly between models with differing dosage compensation specification, as judged by their log likelihoods (P > .05). In the model with equal X-linked variance for males and females, the estimate for the X chromosome variance was 0.34% (SE = 0.07); the variance from the autosomes was 14.37% (SE = 0.42). The model with no dosage compensation (greater female X-linked variance) produced estimates of 0.43% (SE = 0.09) for X-linked variance, and 14.36% (SE = 0.42) for autosomal variance. The model with full dosage compensation (greater male X-linked variance) gave variance estimates of 0.22% (SE = 0.05) for X chromosome, and 14.39% (SE = 0.43) for the autosomes. X-linked variance was significant for all models (P < 6.43×10^−8^).

X chromosome polygenic scores were calculated in unrelated individuals from Generation Scotland ^29^ separately for men and women based on the sex-stratified association analyses. These were used to predict Eysenck Personality Questionnaire neuroticism scores, General Health Questionnaire 28-item ^33^ total scores (i.e., general psychological distress), and Structured Clinical Interview for DSM-IV (SCID)-diagnosed depressive disorder ^34^. Following FDR correction, none of the results were significant (P > .05; Supplementary Table 5).

Because *HS6ST2* was the only gene within which single SNPs attained significance, was mapped within the key region at Xp26.2, and had been implicated in a relevant behavioural trait in the dog ^35^, it was investigated further. A phylogenetic tree was constructed from all available orthologous sequences (Supplementary Figure 12) and showed that the dog sequence was in a group with the cat and leopard, and close to the horse, another companion animal, and to the alpaca, a domesticated species. Somewhat more distant were other domesticated species—the goat, pig and sheep—but these were noted to have fragmented and incomplete sequences which could affect the analysis. In all mammalian species where a chromosome had been assigned, the orthologous gene was on the X chromosome; where there was no chromosome assigned, the gene was in a region syntenic with the X chromosome region of human and was flanked by the same set of genes (*MBNL3*, *RAP2C* and *FRMD7* on one side and *USP26*, *GPC4* and *GPC3* on the other). Alignment of the *HS6ST2* sequence from a selection of mammals, birds and fish revealed that there was strong conservation of the amino acid sequence across the protein, particularly within the putative transmembrane domain, the P-loop containing NTP hydrolase domain, and the region omitted by alternative splicing (highlighted in yellow, green and pink respectively in Supplementary Figure 13). The dog and human predicted protein and cDNA sequences for HS6ST2 showed 93% identity (Figure 2), whereas the similarity between dog, cat and horse was 96% to 98% (Supplementary Figure 13). In contrast, for another X chromosome gene, *HPRT1*, which maps within 2 mbp of *HS6ST2* in most mammals, dog and human showed 100% identity whereas cat showed 93% identity with human/dog and 89% with horse. The dN/dS ratio between dog and human was 0.17, indicating that there has been strong selection pressure to conserve the amino acid sequence ^36,37^; this is consistent with the alignment of all species (Supplementary Figure 13).

## Discussion

We have identified three independent loci on the X chromosome associated with scores on the personality trait of neuroticism. In gene-based tests, 7 genes were associated with neuroticism. We could not discriminate between different gene dosage compensation models, although X chromosome variance was significant in each one. Assuming equal X-linked variance between the sexes, we found that 0.34% of variance in neuroticism was explained by the X chromosome, which represents 2.3% of the total variance due to common genetic variants. A composite score of these variants did not predict variance in neuroticism and related traits in an independent sample. The XY pseudoautosomal region did not uncover any significant variants.

Significant SNPs were mapped to a protein coding gene, *HS6ST2*; this gene was also the most significant one arising from the gene-based tests. This gene encodes a member of the heparan sulfate sulfotransferase gene family and has a role in cranial nerve development ^38^. Two isoforms identified by cDNA cloning in humans showed the long form to be expressed in brain-related tissue ^39^. Sociability traits (attitude to human strangers) have been previously mapped to *HS6ST2* in dogs ^35^, with the neurobiological pathways involved in these emotions highly conserved in vertebrates ^40,41^. We found that the amino acid and nucleotide sequences are highly conserved between dogs and humans, particularly in the transmembrane and NTP hydrolase domains. The canine sequence was even more similar to other companion animals, the cat and horse. Since the similarity for another X chromosome gene, *HPRT1*, was different, it is possible that this reflects selection for trait(s) associated with this gene. Sociability would have been strongly selected for in the evolution of companion animals, which may explain the strong conservation of this sequence. The high degree of conservation across vertebrates shown by the phylogenetic tree and the dN/dS ratios indicate that this gene encodes a protein which is under strong evolutionary constraint, particularly in the catalytic domain.

Six additional genes were found to be significant in the gene-based tests. *IL1RAPL1* (Interleukin 1 Receptor Accessory Protein Like 1) is involved in neurite outgrowth, synapse formation and stabilisation. Deletions and mutations in this gene have been associated with intellectual disability and autism spectrum disorder ^42,43^. Further, there has been consistent support for a link between methylation of *IL1RAPL1* and major depressive disorder (MDD), with identified probes showing higher methylation in MDD cases than controls ^42,44^. *PHF6* (Plant Homeodomain (PHD)-like Finger protein 6) mutations are associated with Börjeson-Forssman-Lehmann syndrome with some documented cases characterised by behavioural disturbance related to anxiety, compulsive behaviour, and emotional attachment ^45^. A variant close to *NLGN4X* (neuroligin 4 X-linked) has been associated (at a nominal level) with a quantitative index of suicidality in 2,023 depressive cases ^46^. Mutations in this gene have been associated with autism spectrum disorder ^47^ and a role of neuroligins in cognitive function has been theorised to occur through cell adhesion and synapse formation pathways 48.

Our analysis has identified a number of X chromosome genes that are promising to follow up in future analysis. The increase in power from a previous study of neuroticism that did not find significant X chromosome SNP heritability ^14^ has enabled us to detect this here. The amount of explained variance is low, only 2.3% of the total common SNP heritability, whereas for physical traits like male pattern baldness, almost 9% of common SNP heritability can be explained by the X chromosome ^49^. Given this low variance, it is unsurprising that polygenic scores did not predict neuroticism in an independent sample. Polygenic scores created from autosomes explain up to 3% of variance in neuroticism ^3^, but the heritability due to common autosomal SNPs is 14% compared to about 0.3% for X chromosome variants. Further, polygenic scores were created from the sex-stratified association analyses which had less power than the meta-analysis; with larger sample sizes, X chromosome polygenic prediction may improve.

In UK Biobank, we have found that common SNPs on the X chromosome contribute to the heritability of neuroticism and we identified SNPs and genes that were associated with this important personality trait. Given the strong autosomal genetic correlations between neuroticism and psychiatric traits ^3^, this study suggests that X chromosome analysis for related traits would successfully identify further genes/biological pathways involved in their aetiology.

## Acknowledgements

This research has been conducted using the UK Biobank Resource (Application Nos. 10279 and 4844). This work was supported by The University of Edinburgh Centre for Cognitive Ageing and Cognitive Epidemiology funded by the Biotechnology and Biological Sciences Research Council (BBSRC) and Medical Research Council (MRC) (MR/K026992/1), which also supports I. J. D, G. D., C. R. G., and D. C. L. W.D.H. is supported by a grant from Age UK (Disconnected Mind Project). A.M.M. is supported by funding from a Wellcome Trust Strategic Award (104036/Z/14/Z). K.M.S. is supported by the Mater Foundation, Brisbane, Australia.

## Data Availability

The X chromosome association results generated by this analysis can be downloaded from the CCACE data sharing resource; **http://www.ccace.ed.ac.uk/node/335**.

## URLs

UK Biobank Resource: http://www.ukbiobank.ac.uk

Gene Ontology: http://geneontology.org

METAL: http://csg.sph.umich.edu/abecasis/metal/index.html

PLINK V2: https://www.cog-genomics.org/plink2

Ensembl: http://www.ensembl.org

Clustal Omega: https://www.ebi.ac.uk/Tools/msa/clustalo/

## Supplementary

